# STAnalyzer: Transparent Spatial Transcriptomics Analysis via an Agentic Architecture

**DOI:** 10.64898/2026.04.06.716827

**Authors:** Hanwen Luo, Liliang Liu, Zhongyu Xing, Xinlu Li, Xuerui Zhang, Wei Du, Bin Liu, Jun Wang, Guoxian Yu

**Author notes:** To whom correspondence should be addressed. Correspondence may also be addressed to Jun Wang and Bin Liu. and. These authors contributed equally to this work (Hanwen Luo, Liliang Liu).

## Abstract

Spatial transcriptomics enables high-resolution profiling of gene expression within spatial contexts, yet its potential is often hindered by fragmented toolchains, intricate parameters, and cognitive bottlenecks of interpreting high-dimensional data. While recent Large Language Model agents have attempted to automate this process, they remain constrained by rigid execution logic, lack multimodal feedback for self-correction, and operate in epistemic isolation from established biological knowledge. Here, we present STAnalyzer, an intelligent multi-agent framework designed to automate the end-to-end analytical lifecycle—from raw data processing to biological hypothesis generation. Transcending traditional pipelines, STAnalyzer employs a collaborative intelligence architecture to achieve three core capabilities: (1) **Intent-Driven Orchestration**, which dynamically translates natural language queries into rigorous bioinformatics workflows; (2) **Multi-Modal Self-Refinement**, which autonomously ensures analytical robustness through closed-loop synthesis of evidence from visual patterns and statistical metrics; and (3) **Evidence-based Cross-Validation**, which bridges the gap between data-driven correlations and biological causation by anchoring findings in ground-truth literature and structured databases. By eliminating manual analytical bottlenecks and ensuring rigorous evidentiary traceability and transparency, STAnalyzer makes high-resolution spatial omics more accessible to a broader research community. It provides a robust and scalable framework for cross-platform automated analysis and accelerated biological discovery, translating massive spatial datasets into verifiable biological insights.

## 1 Introduction

Spatial transcriptomics (ST) has emerged as a transformative technology in modern biology and medicine, enabling the simultaneous capture of gene expression profiles and their spatial contextualization within intact tissues [1, 2]. This unique capability has unlocked unprecedented insights into tissue architecture, cellular crosstalk, developmental dynamics, and disease microenvironment heterogeneity, revolutionizing fields ranging from basic developmental biology to precision oncology and neuroscience [3, 4]. With the rapid advancement of ST platforms (e.g., Visium, Slide-seq [5], and MERFISH [6]), the volume of spatial transcriptomic data has grown exponentially, driving an urgent demand for efficient, standardized, and user-friendly analytical tools [7].

While platforms like SpatialRef [8] and SODB [9] provide essential ST resources, their practical utility is hindered by dual technical and cognitive bottlenecks. Computationally, fragmented pipelines and complex deployments present significant challenges for biologists lacking advanced computational expertise [10, 11, 12]. Biologically, deciphering high-dimensional outputs into actionable insights demands substantial domain knowledge, imposing a considerable cognitive load. Furthermore, current isolated tools intrinsically lack cross-step, cross-result, and context-aware interpretive capabilities. Consequently, this limited systemic integration, coupled with steep knowledge prerequisites, constrains the broader application of ST technologies.

In recent years, the breakthrough progress of Large Language Models (LLMs) in zero-shot learning and generalization has significantly advanced bioinformatics and automated experimental data analysis, inspiring representative works such as Cell Agent [13], Bio Agents [14], and Bioinformatics Agents [15] to build intelligent agents for complex task automation. However, applying these general-purpose bio-agents to spatial transcriptomics data reveals critical deficiencies in handling real-world scientific complexity. First, existing methods often suffer from fragile execution without dependency awareness, as they treat bioinformatics tools as isolated entities while neglecting the implicit logical dependencies between analytical steps and the volatility of fragmented ecosystems [11, 16]; Second, current systems are characterized by open-loop execution with multimodal blindness, operating as “fire-and-forget” pipelines that lack the semantic capability to interpret multimodal outputs [17]. Without a structured strategy to navigate complex results, these agents frequently succumb to context overflow and fail to perform essential self-reflection or dynamic replanning based on intermediate biological signals. Third, these agents suffer from epistemic isolation from external knowledge because they rely on static internal parameters, which prevents them from synergizing structured databases with unstructured literature. Lacking mechanisms for query alignment or coarse-to-fine reading, they cannot retrieve high-fidelity evidence, rendering them unable to provide the traceable, citation-backed insights required to distinguish genuine biological discoveries from technical artifacts [18].

To bridge these critical gaps, we present STAnalyzer, an intelligent multi-agent collaborative framework designed to automate the spatial transcriptomics analysis lifecycle through natural language interaction. By simply providing raw data and a research query (e.g., “Identify genes driving the tumor microenvironment changes and verify with literature”), users trigger an autonomous process that integrates three core capabilities. First, the system excels in autonomous planning and orchestration, decomposing high-level intents into executable bioinformatics workflows that range from data normalization to the generation of multimodal visualizations. Second, STAnalyzer employs self-refining quality assurance through a closed-loop feedback mechanism, critically evaluating intermediate results against analytical standards and autonomously rectifying parameters to ensure robust outcomes. Third, the framework performs evidence-based cross-validation via a dual-pipeline knowledge engine that cross-references computational findings with external ground truth from structured databases and high-impact literature. This process generates traceable, citation-backed insights that bridge the gap between data-driven inference and established biological causation. By automating and enhancing the entire analytical pipeline, STAnalyzer effectively lowers technical barriers, ensures process traceability, and substantially reduces the cognitive load required to translate high-dimensional outcomes into meaningful biological discoveries. Through applications on both spot-level and subcellular-resolution spatial transcriptomic datasets encompassing human brain and lung cancer, STAnalyzer demonstrates its effectiveness in two complementary dimensions: rapidly and autonomously recapitulating established biological paradigms, and serving as a scalable engine for cross-platform automated analysis and accelerated biological discovery.

## 2 Materials and methods

### 2.1 STAnalyzer Architecture: A Human-in-the-Loop Multi-Agent Framework

STAnalyzer is an intelligent collaborative system tailored for comprehensive spatial transcriptomics (ST) data analysis. In contrast to traditional “black-box” automated pipelines, STAnalyzer implements a **Human-in-the-Loop (HITL)** paradigm, seamlessly integrating multiagent autonomy with human domain expertise. The core engine is orchestrated through four functionally specialized agents: the **Orchestrator Agent** (OA), **Service Planner Agent** (SPA), **Data Interpretation Agent** (DIA), and **Knowledge Integration Agent** (KIA). These agents autonomously decompose complex biological workflows, manage execution states, and query external knowledge. However, rather than operating in isolation, the framework embeds the user at the center of the decision-making process, ensuring that computational autonomy is continuously guided, verified, and refined by human intuition.

### 2.2 Interactive Human-in-the-Loop (HITL) Interface

This synergistic architecture is operationalized via a web-based dashboard that establishes a continuous human–AI collaborative loop. Within this interface, the analytical workflow is represented as a dynamic provenance graph. Researchers can select specific intermediate data nodes from this graph and integrate them into the conversational context. While natural language prompts can trigger the agents to autonomously parse requirements and execute bioinformatics workflows, the system strictly maintains process transparency. When granular intervention is required, researchers can override agent autonomy to manually configure execution parameters, define analytical boundaries, and initiate specific services. As intermediate outputs are generated, they can be dynamically reassigned as new context, enabling an iterative, query-driven exploration of the data.

To bridge computational results with biological interpretation, the interface integrates essential spatial visualization capabilities. This allows users to inspect key spatial characteristics, such as cell distribution patterns, gene expression heterogeneity, and spatial co-localization, directly within the analytical context. By cross-referencing these visual cues with agent-generated structured reports and retrieved literature, users can critically evaluate the validity of the computational findings. Ultimately, this mechanism establishes a rigorous analytical loop: system output triggers human verification, which in turn guides subsequent computational interventions and optimizations. If the results deviate from biological expectations, the workflow can be seamlessly rolled back to previous decision nodes for re-evaluation. Ultimately, this HITL design culminates in the delivery of a comprehensive and fully traceable analytical report.

### 2.3 Orchestrator Agent (OA): User-Centric Coordinator

As the sole user-facing entry point and central coordinator of STAnalyzer, the OA functions as a user proxy responsible for translating unstructured natural language requests into structured, executable actions while maintaining global analytical consistency. Upon parsing a user’s research intent, the OA iteratively manages the system by mapping high-level goals to concrete bioinformatics tasks—such as data normalization or spatial domain identification—via explicit service execution directives. Beyond mere task delegation, it actively evaluates execution outcomes and intermediate data files through result inspection and feedback parsing, determining whether current analytical objectives are met or require parameter rectification. This technical workflow is further enriched by knowledge retrieval and integration, which triggers KIA queries to bridge the gap between data statistics and biological significance using external databases and literature. Throughout this lifecycle, the OA anchors the process within a dynamic Global Context Memory—logging project background, data states, and historical interactions—to ensure that every analytical step remains tightly aligned with the user’s scientific hypothesis until a comprehensive, biologically grounded result is achieved.

Throughout this process, the OA maintains a dynamic Global Context Memory, which logs project background, uploaded data states, and historical interaction records. By continuously cycling, inspecting results, and querying knowledge, the Orchestrator ensures that the analysis not only progresses technically but also aligns tightly with the user’s scientific hypothesis until a comprehensive, biologically meaningful final result is generated.

### 2.4 Service Planner Agent (SPA): Addressing Bioinformatics Complexity in Tool Selection and Execution Robustness

SPA is designed to translate abstract research intents into concrete, reliable analytical steps, but its application in bioinformatics is hindered by significant challenges. Specifically, the agent must navigate a vast tool selection space complicated by implicit task dependencies, manage intrinsically unstable execution environments prone to complex programming language and library compatibility issues, and overcome cryptic diagnostic feedback that severely limits its ability to devise effective error recovery strategies. To systematically resolve these interconnected planning, execution, and debugging bottlenecks, we propose a three-layer framework that integrates constraint matching, a robust execution loop, and containerized microservices to ensure seamless and automated analysis.

To systematically address these challenges, we propose a three-layered framework (Figure 2), integrating constraint-based matching, a robust execution loop, and containerized microservices.

**Figure 1:**
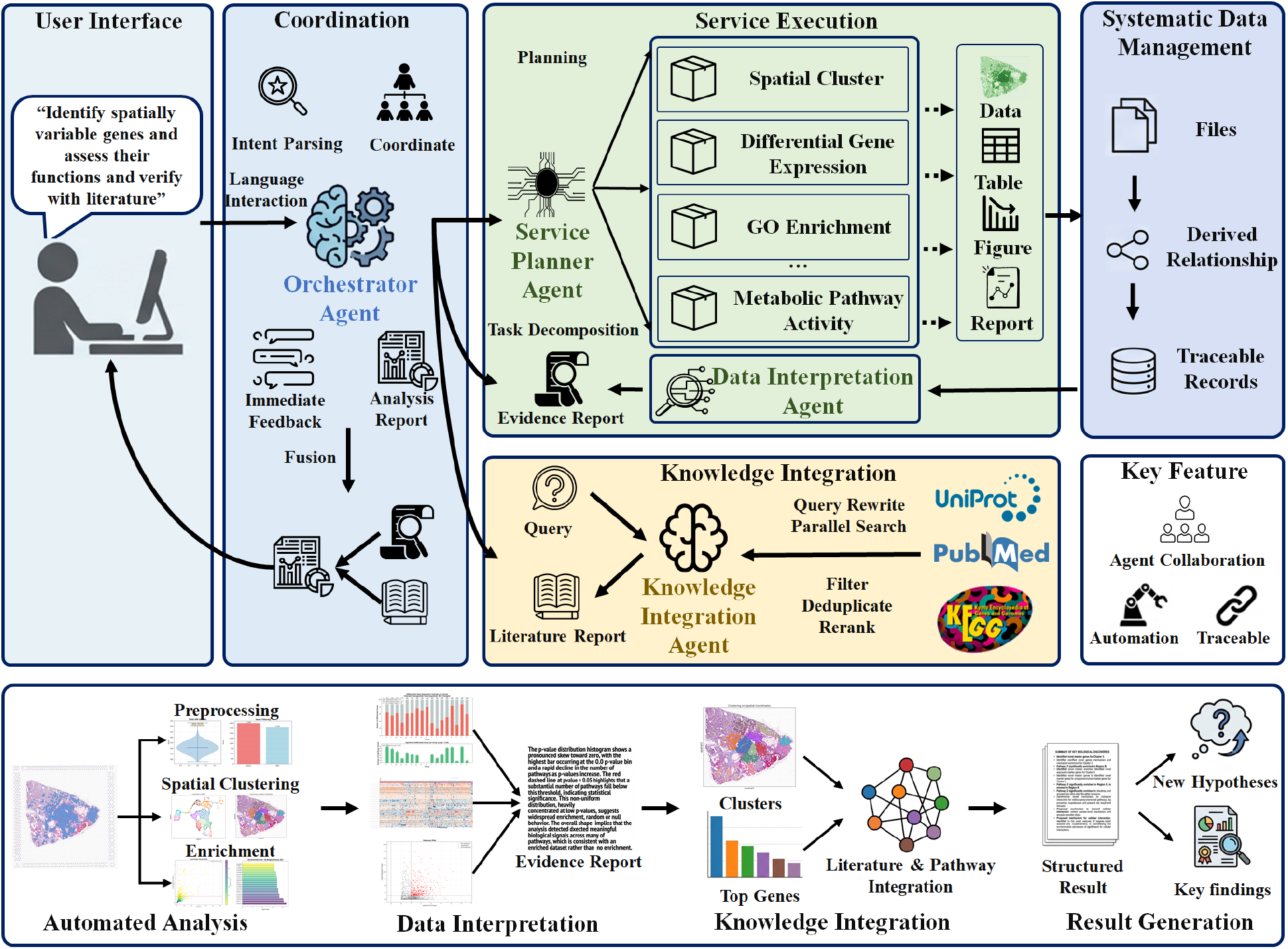
System architecture and end-to-end workflow of STAnalyzer. The framework comprises: (1) a natural language User Interface; (2) the Orchestrator Agent for intent parsing and multi-agent coordination; (3) the Service Execution layer including the Service Planner Agent, specialized analytical services, and the Data Interpretation Agent; (4) the Knowledge Integration Agent linking external resources (PubMed, KEGG); (5) the Data Management module for file provenance and traceability; and (6) an end-to-end workflow from automated analysis through knowledge integration to result generation.

**Figure 2:**
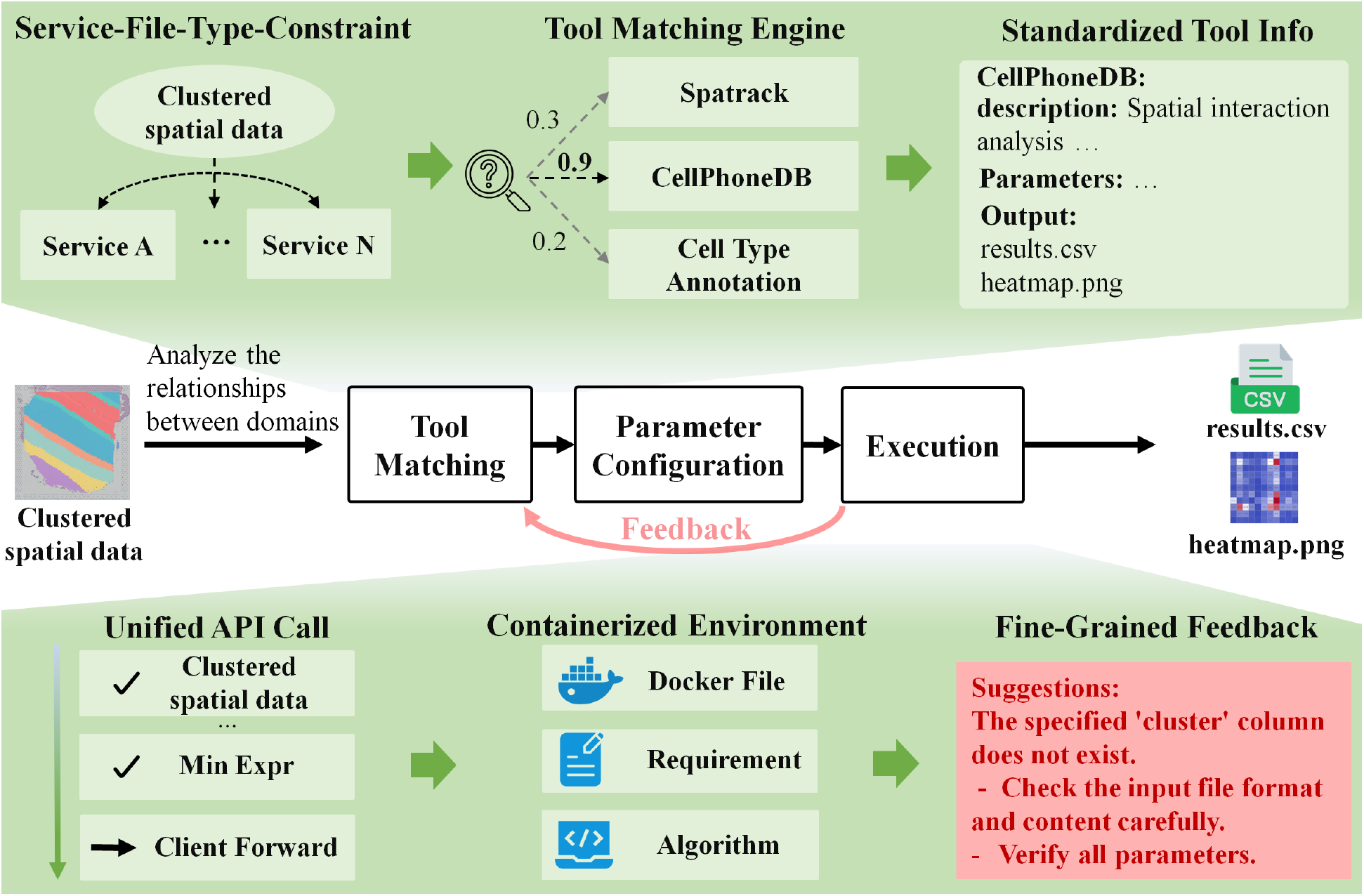
Architecture of the Service Planner Agent. The framework is organized into three logical layers: (Top) The **Tool Matching Layer** leverages Service-File-Type Constraints and a semantic Tool Matching Engine to select appropriate tools (e.g., CellPhoneDB) based on standardized descriptions. (Middle) The **Execution Workflow Layer** illustrates the closed-loop process from tool matching and parameter configuration to execution, incorporating a feedback mechanism for error recovery. (Bottom) The **Infrastructure Layer** supports the workflow via a Unified API Call interface and a Containerized Runtime Environment, providing Fine-Grained Diagnostic Feedback to resolve execution failures.

#### Constraint-based Tool Matching (Top Layer)

To navigate the expansive toolset, we establish a Service-File-Type Constraint mechanism. As shown in the top panel of Figure 2, this mechanism first constrains the search space based on the input data type (e.g., mapping “Clustered spatial data” to candidate tools like spatrack and CellPhoneDB). Subsequently, the Tool Matching Engine evaluates these candidates against the user’s specific intent. By comparing the intent with **Standardized Tool Info**, the engine assigns relevance probabilities—for instance, assigning a high confidence score of 0.9 to CellPhoneDB for interaction analysis tasks, while assigning lower scores (e.g., 0.2 or 0.3) to less relevant tools.

#### Robust Execution Workflow (Middle Layer)

The middle panel of Figure 2 illustrates the agent’s robust execution workflow, which operates as a closed loop explicitly driven by both data and research intent. Upon receiving input data, the agent first utilizes a Matching Engine to select the optimal analytical tool and configures its parameters based on standardized metadata. The task is then dispatched through a unified API call, ensuring standardized interaction regardless of the underlying programming language, such as R or Python. Following execution, the system either generates successful analytical results or, in the event of a failure, activates a continuous feedback mechanism that captures fine-grained diagnostic information. This feedback loop empowers the agent to iteratively adjust parameters or re-plan the analysis, significantly enhancing its overall robustness against runtime errors.

#### Containerized Microservices with Fine-Grained Feedback (Bottom Layer)

To resolve environment fragility, we refactored bioinformatics tools into a Microservices Architecture. As illustrated in the bottom panel of Figure 2, tasks are dispatched via unified API call, which encapsulates the data and parameters (e.g., `Min expr`). Each tool runs within an isolated containerized runtime environment (defined by Dockerfile and Requirements). Crucially, the system replaces cryptic stack traces with fine-grained feedback. For example, when a “cluster label column not found” error occurs, the system generates structured suggestions (e.g., “Check input file format”, “Verify parameters”), enabling the agent to pinpoint the root cause (e.g., column name mismatch) rather than blindly retrying.

Furthermore, the agent incorporates mechanisms to identify similar failure patterns, ensuring that non-productive execution cycles are terminated after repeated failures, redirecting the process to result reporting, thus guaranteeing system stability and reliability. For customized needs beyond the scope of predefined tools, SPA also supports generating and executing customized analysis scripts in a secure isolated environment, integrating the results into the overall analytical sequence.

### 2.5 Data Interpretation Agent (DIA): Multi-Modal Understanding and Synthesis

DIA is designed to interpret computational outputs during spatial transcriptomics analysis. As illustrated in **Figure 3**, the agent operates in two distinct modes to accommodate different analytical granularities: **(A) Single File Query** for specific data inspection, and **(B) File Tree Query** for complex, multi-file synthesis.

**Figure 3:**
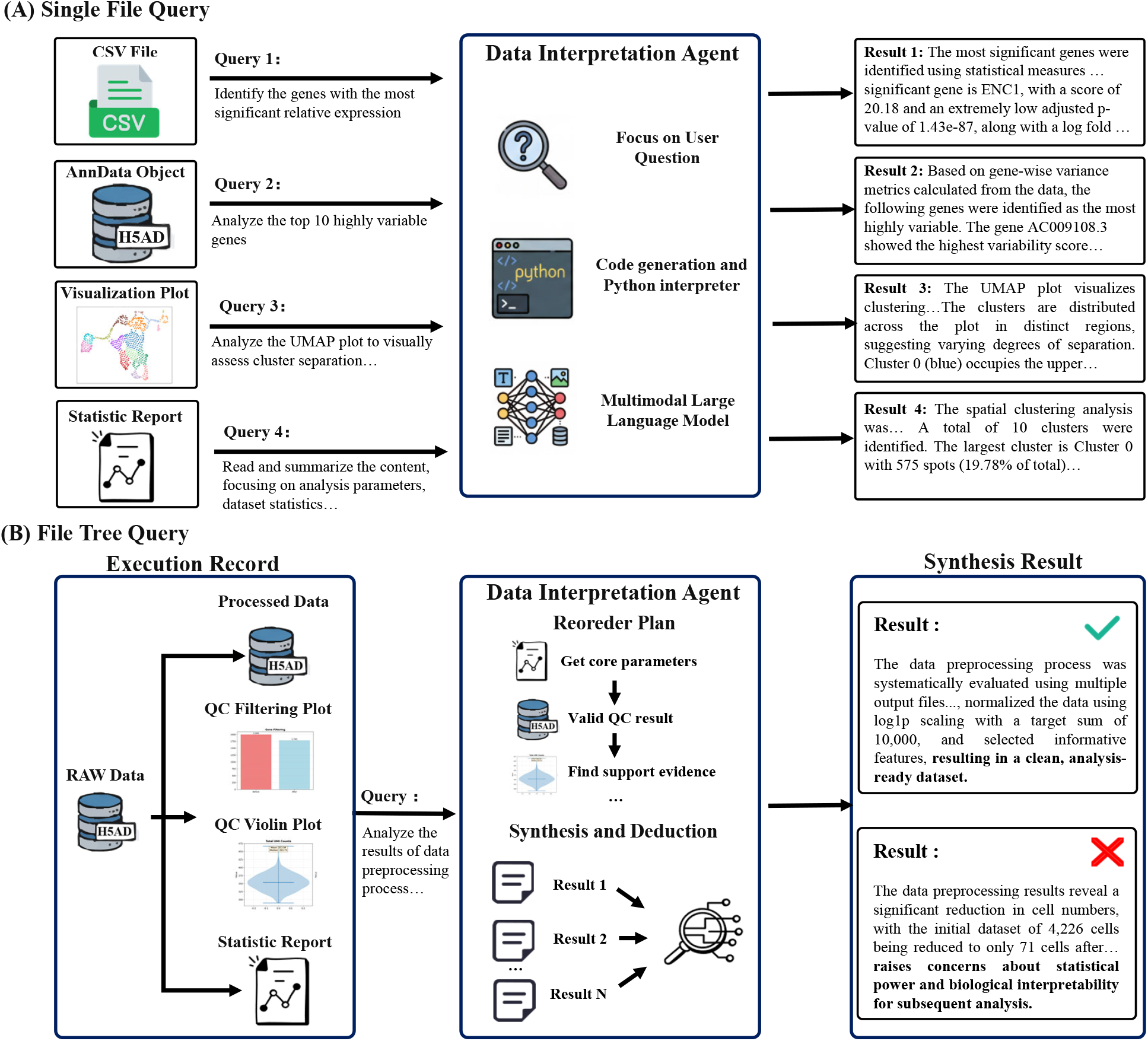
Workflow of the Data Interpretation Agent. **(A) Single File Query:** The agent processes individual files (CSV, H5AD, Images, Reports, etc.) based on specific user queries. It utilizes tools like code interpreters and multimodal models to extract gene signatures, statistical metrics, and visual cluster patterns. **(B) File Tree Query:** For complex synthesis tasks, the agent inputs the entire execution record. It employs a “Reorder Plan” to prioritize data access (e.g., getting parameters before evidence), performs synthesis and deduction, and outputs a final validation result (Pass/Fail) regarding the analysis quality.

#### Single File Query Mode

As shown in **Figure 3(A)**, the agent possesses multi-modal capabilities to handle diverse file formats individually. It accepts user queries targeting specific outputs, such as identifying significant genes from CSV files, analyzing variable genes in AnnData objects (H5AD), or interpreting visualization plots (e.g., UMAP, spatial maps). By leveraging code generation tools and Multi-modal Large Language Models (MLLMs), the agent extracts statistical metrics and visual features to provide precise, isolated answers.

#### File Tree Query Mode

This mode is designed for comprehensive assessments that require synthesizing information across an entire execution record. Instead of indiscriminately reading all files, which risks context overflow, the agent employs a dynamic **Reorder Plan** to logically sequence data access based on the query’s intent. As illustrated by the Quality Control (QC) validation example in **Figure 3(B)**, this plan prioritizes the workflow: first extracting core parameters, then validating numerical results, and finally seeking supporting evidence from plots (e.g., Violin plots). This structured approach enables the agent to aggregate fragmented evidences into a cohesive synthesis, ensuring that the final conclusion is both biologically interpretable and statistically robust.

To ensure analytical accuracy across both modes, the agent applies tailored logic to seamlessly handle diverse file types. When processing text reports, it targets the extraction of core conclusions and experimental design parameters, condensing the content to focus strictly on key analytical indicators. For structured data, such as CSV or AnnData files, the agent couples file previews with automated Python code generation to query the data, effectively translating complex numerical matrices into human-readable insights on gene expression and variability. Furthermore, when analyzing image files like plots, it leverages advanced vision-language capabilities to objectively interpret visual patterns—such as UMAP cluster separation or distributions in QC plots. By aligning these visual interpretations with numerical data for rigorous cross-validation, the agent accurately delineates biological implications while strictly preventing hallucinations. Ultimately, this unified approach enables the agent to synthesize text, numerical matrices, and images, delivering a comprehensive analysis of heterogeneous data.

This design, centered on ordered reading plans, effectively enables multi-file joint analysis while maintaining system stability. It ensures that the agent’s contextual reasoning remains precise, reduces the cognitive burden on users, and delivers interpretable, high-quality results that directly support biological hypothesis generation and validation.

### 2.6 Knowledge Integration Agent (KIA): Contextualization of Results with External Biological Knowledge

KIA is designed to resolve a critical bottleneck in spatial transcriptomics analysis: the interpretation of computational results (e.g., region-specific gene expression patterns) is often constrained by a lack of external biological context. Unlike conventional tools limited to isolated analytical findings, KIA automates the retrieval, synthesis, and cross-validation of high-relevance external knowledge, thereby streamlining the validation of biological plausibility without requiring manual literature synthesis.

Biological knowledge sources exhibit inherent heterogeneity and complementary utility. Structured biological databases (e.g., CellMarker, BioGrid, KEGG, NichNet) provide relatively precise and stable conclusions regarding molecular interactions, though they may occasionally suffer from curation lags. In contrast, academic literature offers timely experimental evidence, specific contextual constraints, and cutting-edge insights. However, the utility of literature is often challenged by its massive volume and unstructured nature. The vast quantity of textual data, combined with high information density, makes rapid information extraction and systematic reading computationally demanding for human researchers.

The workflow of KIA is driven by two parallel pipelines initiated by a query alignment module. Biomedical queries typically involve complex semantics with multiple entities (e.g., genes, pathways, diseases) and constraints (e.g., species, experimental conditions). To address this, KIA first performs query alignment to parse the semantic context, extracting core entities and constraints. The original query is then rewritten into a standardized format. This preprocessing ensures retrieval accuracy and mitigates deviations caused by ambiguous semantics.

The first pipeline targets unstructured literature data by leveraging advanced Retrieval-Augmented Generation (RAG) technology [19]. The rewritten query is optimized into PubMed advanced search syntax to narrow the search scope within the massive literature pool. Following initial retrieval, a reranker model performs relevance sorting to filter noise, retaining the top 50 documents in a coarse-ranking stage. To efficiently tackle the challenge of reading unstructured text, KIA generates a targeted *reading plan*. This plan selects the 5 most critical documents and defines specific extraction objectives (e.g., gene-pathway interaction evidence, experimental methods) for each. This strategy transforms the broad scanning of unstructured text into a focused extraction of high-value evidence, ensuring systematic cross-validation. The second pipeline focuses on structured data, integrating multi-source biological databases. Upon extracting core entities from the standardized query, KIA autonomously invokes corresponding database interfaces to retrieve structured knowledge (e.g., gene regulatory networks from BioGrid, pathway annotations from KEGG).

Finally, KIA synthesizes knowledge obtained from both pipelines to enhance interpretability. The retrieved information is organized logically: background context *→* supportive evidence (dual-source) *→* supplementary explanations. This structured synthesis explicitly links external knowledge to in-house experimental results, enabling users to rapidly discern connections between spatial transcriptomics findings and existing research, thereby facilitating hypothesis generation. A defining characteristic of KIA is the strict **traceability of biological evidence**. All integrated external knowledge is presented with granular source attribution, including manuscript titles, contextual excerpts, URLs, and Digital Object Identifiers (DOIs). This transparency serves a dual purpose. First, it facilitates human verification, allowing researchers to directly access original sources for rapid fact-checking and citation during manuscript preparation. Second, and crucially, it establishes a verifiable ground truth for inter-agent collaboration. Downstream agents, such as the Orchestrator Agent and Data Interpretation Agent, utilize this evidence as a reference framework to evaluate analytical outcomes—specifically, assessing the concordance between observed spatial expression patterns and reported mechanisms, or identifying potential discrepancies that may spark novel biological findings.

## 3 Results

We evaluated STAnalyzer on two representative spatial transcriptomics datasets spanning different platforms and resolutions to validate its end-to-end analytical capabilities, biological interpretability, and cross-platform scalability: (1) a spot-level 10x Visium human DLPFC dataset (3,673 spots, 33,538 genes) for assessing automated analysis and biological consistency with known cortical architecture, and (2) a subcellular-resolution 10x Xenium human lung cancer dataset (161,000 cells, 480 genes) for demonstrating scalability and the capacity for hypothesis-driven biological discovery.

### 3.1 STAnalyzer enables rapid, biologically interpretable automated analysis of spatial transcriptomics

To demonstrate the comprehensive conversational automated analysis capabilities and practical utility of STAnalyzer for spatial transcriptomics data, we applied the framework to the 10x Genomics Visium human dorsolateral prefrontal cortex (DLPFC) dataset [3]. Comprising 3,673 spots and 33,538 genes, the analysis commenced with the prompt: “Please perform data preprocessing on this slice and briefly describe the quality of the processed data.” STAnalyzer automatically executed preliminary statistical profiling (Fig. 5a) and selected optimal preprocessing strategies to generate quality-controlled data. The processed dataset retained 3,234 spots and 19,877 genes (Fig. 5a). Crucially, the chosen strategies and parameter settings were explicitly displayed in the dialogue interface. The system provided a summary report evaluating preprocessing quality across four dimensions—Robust filtering, Biologically plausible QC profiles, Consistent multi-format validation, and Analysis-ready output, thereby facilitating informed user decision-making. Subsequently, upon receiving the instruction “Please identify the spatial domain of this data and briefly assess the rationality of the resulting spatial domain division” STAnalyzer segmented the tissue into eight spatial domains, revealing a distinct hierarchical architecture (Fig. 5b). Notably, the system provided autonomous evaluation and feedback on the segmentation results. Statistically, it noted that “Domain counts (*n*=8) and size distribution (195–665 spots) reflect biologically interpretable granularity—not overly fragmented nor excessively coarse.” Visually, it confirmed that the “domain umap.png confirms transcriptional separability in low-dimensional embedding, with high intra-domain compactness and only minor, localized inter-domain overlap.” Should the evaluation yield negative feedback, users can iteratively refine and re-execute this step based on the system’s suggestions to ensure biologically meaningful outcomes.

**Figure 4:**
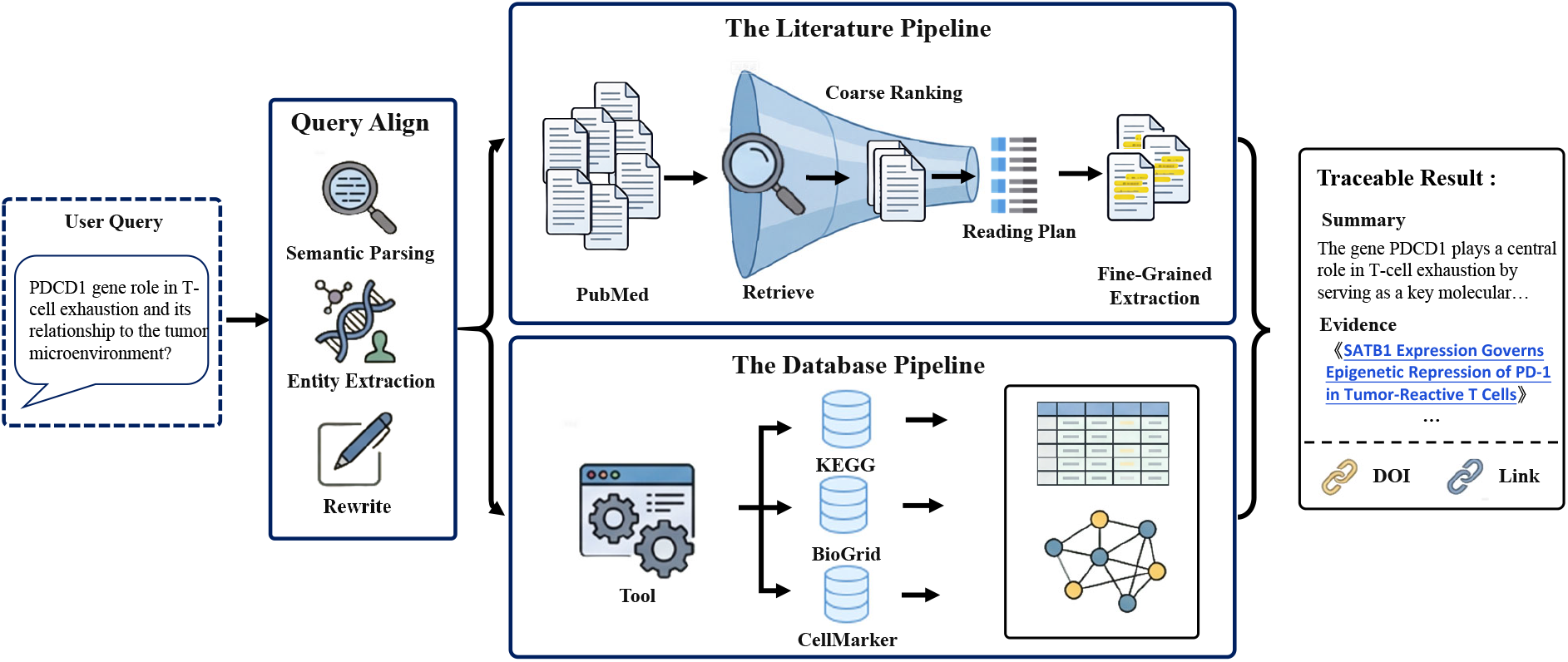
The architecture of Knowledge Integration Agent. The workflow proceeds from left to right, starting with the Query Alignment which standardizes ambiguous user inputs. The process then bifurcates into two complementary pipelines: (Top) The Literature Pipeline employs a RAG-based funnel to process massive unstructured text, using coarse ranking and a targeted reading plan to distill the top 50 candidates down to highly relevant evidence; (Bottom) The Database Pipeline directly retrieves structured molecular facts via unified APIs. Finally, information from both sources is synthesized into Traceable Insights, grounded with explicit citations (DOIs, URLs) to support downstream verification and reasoning.

**Figure 5:**
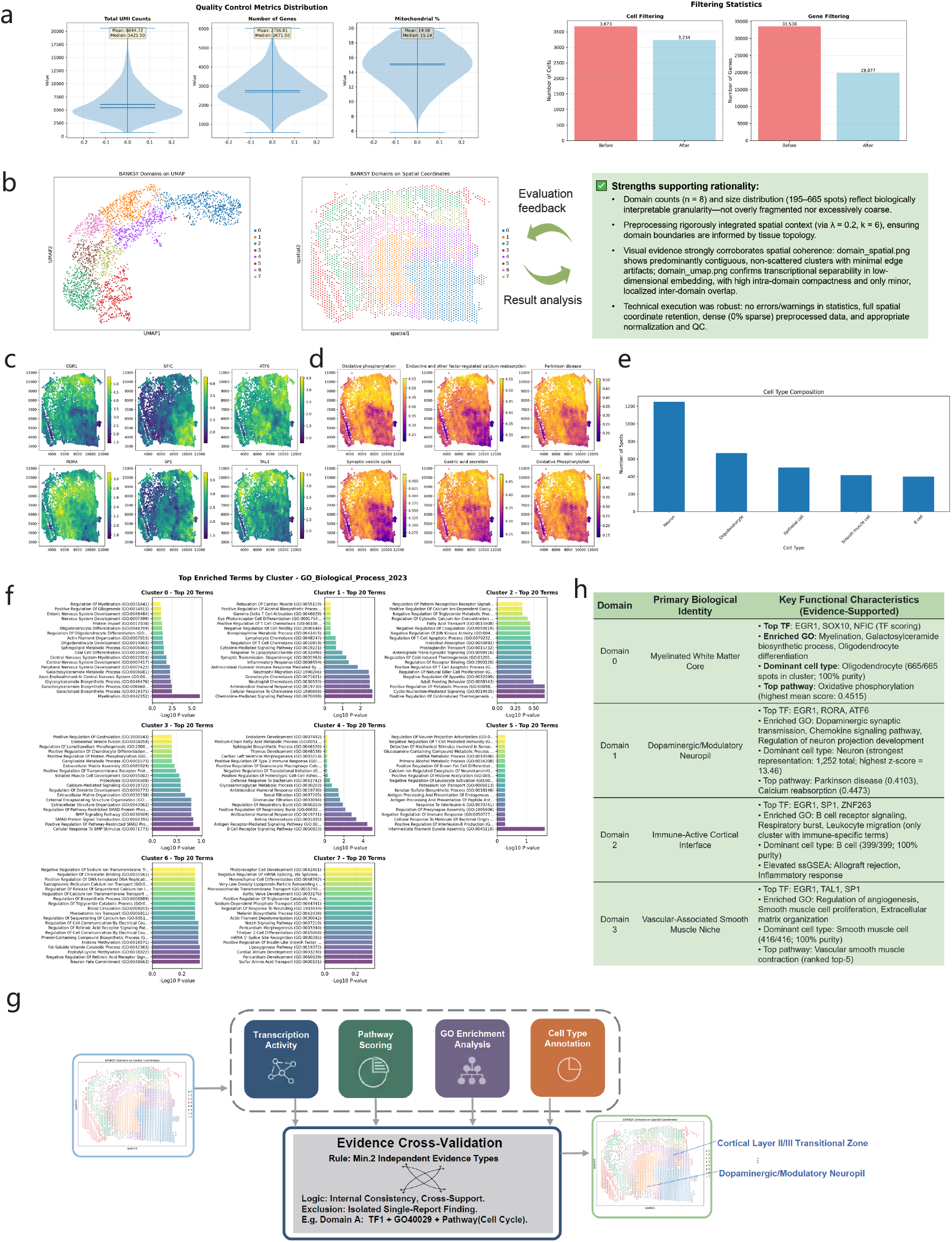
Automated analysis on the human DLPFC dataset. **a** (Left) Statistical analysis of various metrics of the original ST data. (Right) Bar chart of the number of cells and genes before and after quality control. **b** Spatial domain identification results and the automatic assessment feedback provided by STAnalyzer. **c** Heatmap of the top-6 TFs with high spatial TF activity scores. **d** Heatmap visualizing the spatial distribution of major enriched metabolic pathway activities across the ST data. **e** Bar plot of cell type composition. The chart displays the number of spatial spots assigned to each identified major cell type across the tissue section. **f** Top 20 enriched GO terms for each of the eight spatial domains. **g** STAnalyzer conducts evidence cross-validation at four levels to automatically analyze the functional characteristics of each spatial domain. **h** Traceable and interpretable functional characterization of four spatial domains, highly consistent with established biological knowledge.

We then proceeded to a comprehensive functional characterization by prompting: “Based on the obtained 8 spatial domain division results, execute the relevant services to analyze the functional characteristics of each domain. Finally, please provide a table summarizing what each domain is and its corresponding functional characteristics”. STAnalyzer autonomously orchestrated a comprehensive multi-dimensional analysis pipeline. First, it performed Spatial TF Activity Scoring and spatial metabolic pathway activity analysis for each domain. Fig. 5c,d illustrate the top-ranked genes and pathways, highlighting the robust activity of brain-enriched transcription factors (e.g., EGR1, NFIC, ATF6, RORA, SP1, and TAL1) and the significant enrichment of pathways associated with energy metabolism, synaptic transmission, and dopaminergic signaling. These findings closely align with the established physiological and metabolic profiles of the human DLPFC. For instance, EGR1 and RORA are well-documented to be pivotal for synaptic plasticity and neuronal survival in the prefrontal cortex [20, 21]. Similarly, the prominent activity in energy supply and dopamine metabolism pathways is a hallmark of the DLPFC, underpinning its complex executive functions and working memory capabilities [22, 23]. Furthermore, cell type annotation and Gene Ontology (GO) enrichment analyses demonstrated biological plausibility, with cell types predominantly identified as neurons and oligodendrocytes [3]. Leveraging this multi-dimensional evidence, STAnalyzer executed an automated evidence cross-validation process (Fig. 5g). Through internal consistency checks and multi-evidence triangulation, it defined the functional identity of each spatial domain. Four domains exhibited precise alignment with known biological annotations of the human prefrontal cortex (Fig. 5h). For instance, Domain 0 was defined as the myelinated white matter core, driven by oligodendrocyte-specific myelination and oxidative phosphorylation [24]. Cortical neuronal niches were captured by Domain 1 and Domain 4, which highlighted dopaminergic synaptic transmission [25] and neurotrophin-guided synaptic plasticity [26] in superficial layers, respectively. Moreover, the identification of a vascular-associated smooth muscle niche (Domain 3) accurately reflected the cerebrovascular architecture essential for cortical homeostasis [27]. Notably, even without any prior tissue-type annotation or predefined domain count—where the system received no task-specific configuration regarding the input data, the multi-agent collaboration within STAnalyzer accurately reconstructed the spatial architecture and functional characteristics of the human brain slice. Moreover, this comprehensive multi-dimensional validation was completed within minutes, providing detailed, traceable evidentiary support that significantly accelerates the research workflow.

Alternatively, users can opt for a fully automated workflow by inputting a high-level directive such as “Please analyze what functional characteristics this data possesses.” In this mode, STAnalyzer decomposes the request into a transparent, executable sequence of tasks (e.g., data preprocessing, spatial domain identification), executes them sequentially, and generates a final report supported by evidence. While this approach offers greater convenience, it involves less human-in-the-loop interaction compared to the semi-automated mode, offering fewer opportunities for intuitive intermediate feedback and iterative user intervention.

### 3.2 STAnalyzer facilitates biological discovery via cross-platform automated analysis at subcellular resolution

To further demonstrate the cross-platform scalability and advanced exploratory capabilities of STAnalyzer, we applied it to a subcellular-resolution 10x Xenium spatial transcriptomics dataset of human lung cancer [28, 29]. Comprising over 161,000 cells and 480 genes, this dataset represents a cellular scale nearly 50-fold larger than that of brain slice spots. Following initial preprocessing (Fig. 6a), automated spatial domain identification was executed, yielding a highly dense and structurally intact pathological tissue representation comprising nine distinct spatial domains (Fig. 6b). Subsequently, we inputted the natural language query: “Please analyze the main cell type composition of each of the 9 spatial domains one by one (Each spatial domain contains more than one type of cell, and it is composed of multiple cell types), and perform gene enrichment analysis for each domain. Finally, summarize the analysis results for each spatial domain in tabular form.” STAnalyzer automatically executed the analysis and generated plots, detailing significantly enriched terms (Fig. 6c) and the primary cell type composition (Fig. 6d) and for each spatial domain.

**Figure 6:**
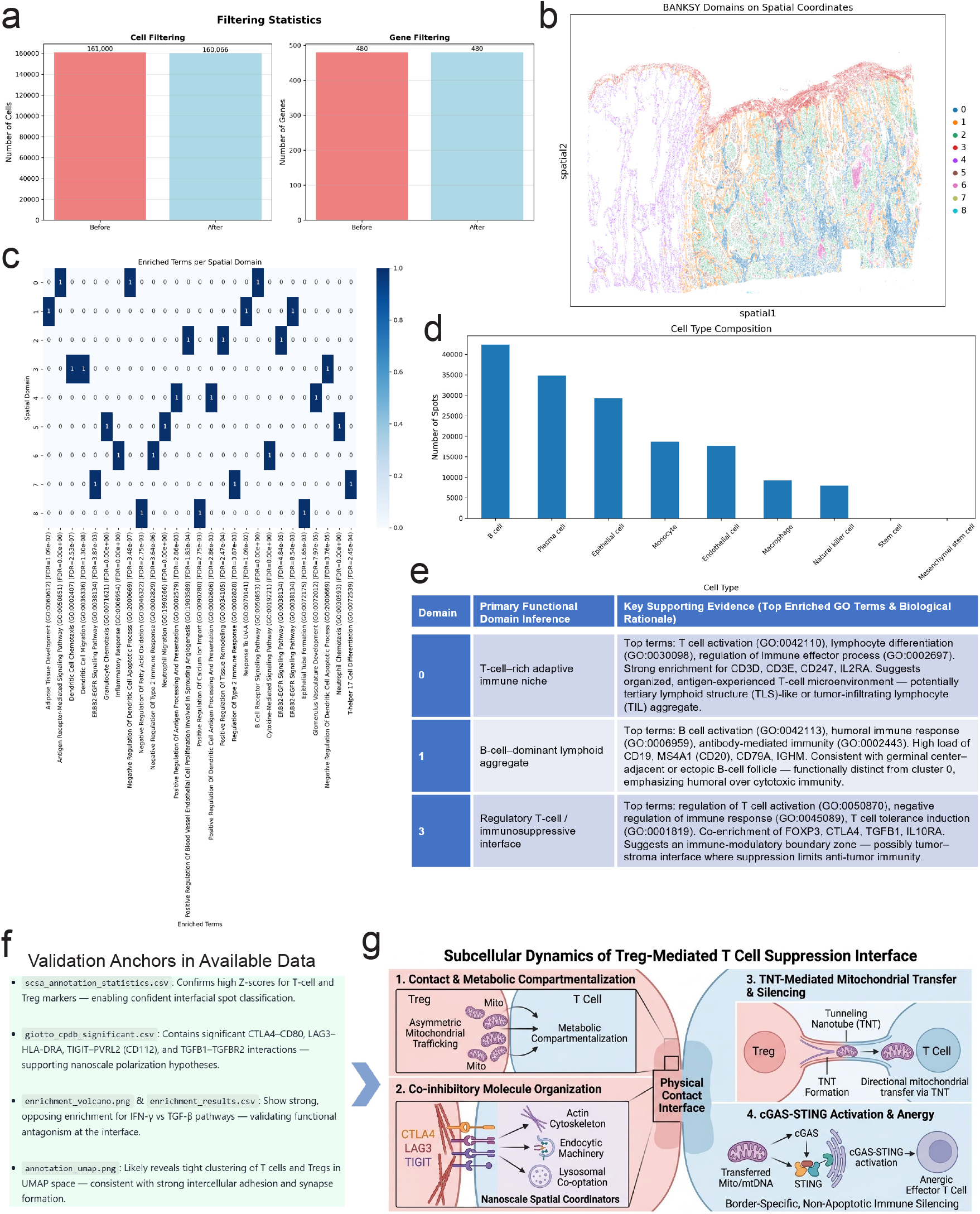
Extended experiments on the 10x Xenium human lung cancer dataset. **a** Bar chart of the number of cells and genes before and after quality control. **b** Spatial domain identification results by STAnalyzer. **c** Heatmap of the top 3 GO terms enriched in each spatial domain. **d** Bar plot of cell type composition. The chart displays the number of spatial spots assigned. **e** The inferred functional results of the main spatial domain and the explainable supporting evidence. **f** STAnalyzer automatically formulates new biological hypotheses based on the available direct evidence. **g** Subcellular dynamics of the Treg-mediated immunosuppressive interface. The physical boundary between Tregs and T cells acts as an active signaling hub. Contact-driven spatial reorganization of co-inhibitory molecules and TNT-mediated mitochondrial transfer synergistically activate the cGAS-STING pathway in T cells. This triggers non-apoptotic immune silencing (anergy), redefining the static barrier as a dynamic target for precision immunomodulation.

Building upon this, we prompted the system to automatically integrate the experimental results for fine-grained functional compartmentalization: “Based on the cell type annotations and gene enrichment analysis results of these 9 spatial domains that have been completed, please further analyze and infer the functional domains that each spatial domain may belong to (provide explainable evidence). Finally, form a summary table to clearly display the characteristics of each spatial domain”. Notably, two compelling phenomena emerged from this analysis. First, STAnalyzer delineated Domain 3 as a continuous, band-like spatial region. Spatial mapping and functional analysis (which showed significant enrichment for the regulation of T cell activation and T cell tolerance induction, alongside high expression of FOXP3 and TGFB1) revealed that Domain 3 constitutes an immunosuppressive physical interface dominated by regulatory T (Treg) cells (Fig. 6e). This intuitively illustrates how tumors spatially construct physical barriers to exclude cytotoxic T cells from the core region [30, 31]. Second, STAnalyzer deconstructed the complex immune infiltration region into two functionally distinct yet spatially adjacent microenvironments: Domain 0 and Domain 1 (Fig. 6b, e). Functional analysis indicated that Domain 0 is enriched for T cell activation pathways (CD3D/E, IL2RA), representing an antigen-experienced T cell microenvironment. Conversely, Domain 1 is highly enriched for B cell activation and humoral immune pathways (CD19, MS4A1). This spatial adjacency and zonation of cytotoxic and humoral immunity strongly suggests the formation of tertiary lymphoid structures (TLS) or ectopic lymphoid follicles [32]. Collectively, these findings demonstrate that STAnalyzer can not only identify cell types but also autonomously decipher complex, higher-order topological structures within the immune spatial microenvironment.

To demonstrate the potential of STAnalyzer in accelerating new, interpretable biological discoveries at the subcellular level through multi-agent collaboration, we provided the following prompt: “The current dataset possesses a truly accurate resolution of subcellular physical coordinates. Our previous analysis has successfully identified a crucial physical communication boundary: Domain 0 (the core of adaptive immunity dominated by T cells) and Domain 3 (the immunosuppressive barrier dominated by Tregs). Please focus on the interface where physical contact occurs between Domain 0 and Domain 3 (the boundary). From the perspective of subcellular analysis, deduce and extract potential new biological discoveries. Please conduct a deep analysis using an explainable thought chain (Chain of Thought).” STAnalyzer autonomously generated code and synthesized all available evidence extracted from file metadata, decision history, and the established knowledge base to propose in-depth, mechanism-based inferences. Crucially, these insights are supported by traceable chains of thought and output files (Fig. 6f), ensuring they are firmly grounded in data rather than being baseless fabrications, thereby significantly mitigating the occurrence of large language model hallucinations [33]. The comprehensive analysis by STAnalyzer concluded: “The Domain 0/Domain 3 interface is not a passive diffusion barrier, but an active, organelle-engineered decision hub, where: (i) Physical contact triggers asymmetric mitochondrial trafficking to enforce metabolic compartmentalization; (ii) CTLA4, LAG3, and TIGIT operate not as isolated receptors, but as nanoscale organizers of endocytic, actin, and lysosomal machinery; (iii) Tunneling nanotubes mediate directional, suppressive mitochondrial transfer, converting T cells into transiently anergic effectors via cGAS–STING — a boundary-specific, non-apoptotic immune silencing mechanism. This reframes the immunosuppressive barrier not as static cellular segregation, but as a dynamic, subcellularly orchestrated interface — offering new targets for precision immunomodulation (e.g., inhibiting TNT formation in autoimmunity, or enhancing mitochondrial transfer in cancer immunotherapy).” Based on the aforementioned subcellular spatial analysis, the immunosuppressive barrier between Domain 0 and Domain 3 is characterized not as static cellular segregation, but as a dynamic, subcellularly orchestrated interface (Fig. 6g). This perspective offers novel targets for precision immunomodulation (e.g., inhibiting TNT formation in autoimmunity, or enhancing mitochondrial transfer in cancer immunotherapy). Notably, these computationally derived hypotheses share complementary parallels with recent experimental evidence [34, 35], further illustrating the capacity of STAnalyzer for cross-platform automated analysis and accelerated biological discovery.

In conclusion, while computational predictions made by STAnalyzer cannot replace rigorous experimental validation as the gold standard for biological discovery, it serves as a valuable tool for generating high-confidence hypotheses. By autonomously executing subcellular spatial transcriptomic analysis of over 100,000 cells within tens of minutes, it transforms massive datasets into evidence-based, interpretable insights. This framework lowers the technical barrier to complex computational research, provides reliable starting points for experimental design, and offers a practical engine for cross-platform automated analysis and accelerated biological discovery.

## 4 Discussion

Spatial transcriptomics has revolutionized our understanding of tissue biology by linking gene expression to spatial context, yet the translation of high-dimensional ST data into actionable biological insights remains hindered by fragmented toolchains, steep technical barriers, and the disconnect between computational results and biological knowledge. In this study, we present STAnalyzer, an intelligent multi-agent framework that addresses these critical gaps through a Human-in-the-Loop (HITL) collaborative architecture, integrating intent-driven orchestration, multi-modal self-refinement, and evidence-based cross-validation. Here, we discuss the core contributions, comparative advantages, limitations, and future directions of STAnalyzer, as well as its broader implications for the field of spatial omics analysis.

Unlike “black-box” pipelines or isolated agents, STAnalyzer’s four specialized agents collaborate to automate end-to-end analysis while retaining human oversight. Key innovations include robust execution via constraint-based tool matching and containerization, semantic integration of multimodal outputs, and dual-pipeline (literature/database) knowledge integration—resolving fragility, multimodal blindness, and epistemic isolation in existing tools. The HITL paradigm balances automation with human intuition, empowering non-computational researchers to leverage ST analysis without sacrificing control or transparency. Practically, STAnalyzer democratizes access to spatial transcriptomics by substantially lowering computational barriers and ensuring reproducibility through traceable, evidence-based reporting. Crucially, the framework exhibits exceptional cross-platform scalability, seamlessly adapting from macroscopic tissue profiling (e.g., analyzing thousands of spots via 10x Visium) to the subcellular-resolution mapping of massive cellular cohorts (e.g., over 100,000 cells via 10x Xenium). By successfully navigating diverse and complex biological contexts—ranging from neuroarchitecture to precision oncology—STAnalyzer drastically reduces manual analytical workloads. Ultimately, it serves as a robust engine for cross-platform automated analysis and accelerated biological discovery, streamlining rapid hypothesis generation and evidence synthesis.

Limitations include a current focus on standard 2D transcriptomic tasks, with future work needed to expand coverage to spatial multi-omics and 3D modalities; reliance on static knowledge sources, which will require real-time synchronization mechanisms; and potential usability gaps for novice users. Future directions involve expanding the tool library, integrating real-time knowledge updates, simplifying user interactions, optimizing distributed computing for ultralarge datasets, and enhancing robustness to noisy data. Ultimately, STAnalyzer represents a paradigm shift toward collaborative intelligence in spatial omics, bridging computational efficiency with biological relevance to drive cross-platform automated analysis and accelerated biological discovery.

As ST technologies continue to evolve, intelligent analytical frameworks will play an increasingly important role in translating spatial data into biological discoveries [36]. STAnalyzer provides a robust, user-friendly solution for end-to-end ST analysis, and ongoing refinements will strengthen its utility as a versatile tool in spatial omics research.

## 5 Webserver Implementation

STAnalyzer is built as a high-performance, scalable web service, designed specifically to support complex multi-agent collaborative workflows and the processing of large-scale ST data. The backend adopts a microservices architecture, containerized with Docker for isolation and scalability, and uses FastAPI on Uvicorn for high-concurrency, low-latency APIs. MongoDB handles structured logs and metadata, while network file systems manage large raw and intermediate data. Intelligent orchestration is powered by LangChain and LangGraph, with Qwen integration for natural language understanding and dynamic decision-making. Tasks are executed asynchronously—each receives a unique Execution ID, and real-time progress is streamed to the frontend via Server-Sent Events. The frontend is a Vue 3-based single-page application using TypeScript and Vite, with Element Plus for UI consistency and Pinia for state management. Interactive workflow visualizations are rendered with AntV G6, and analytical charts are generated using Apache ECharts. Token-based authentication ensures security without requiring manual password entry., and the platform is fully compatible with modern browsers (Chrome, Firefox, Edge, Safari), providing researchers with an interactive interface for data exploration, real-time visualization, and result download.

## 6 Data availability

The STAnalyzer platform is publicly accessible at https://www.sdu-idea.cn/STAnalyzer, with no registration or access restrictions for academic and non-commercial use.

## 7 Funding

This work is supported by National Key Research and Development Program (No. 2023YFF0725500 and 2024YFF1206604) and National Natural Science Foundation of China (No. 62531013, 62432006 and 62272276), Natural Science Foundation of Shandong Province (ZR2024JQ001 and ZR2025LZH004) and Taishan Scholars Program (tsqn202408317).

### 7.0.1 Conflict of interest statement

None declared.

## Notes

### Competing Interest Statement

The authors have declared no competing interest.

## References

[1] Anjali Rao, Dalia Barkley, Gustavo S França, and Itai Yanai. Exploring tissue architecture using spatial transcriptomics. Nature, 596(7871):211–220, 2021.

[2] Vivien Marx. Method of the year: spatially resolved transcriptomics. Nature Methods, 18(1):9–14, 2021.

[3] Kristen R Maynard, Leonardo Collado-Torres, Lukas M Weber, Cedric Uytingco, Brianna K Barry, Stephen R Williams, Joseph L Catallini, Matthew N Tran, Zachary Besich, Madhavi Tippani, et al. Transcriptome-scale spatial gene expression in the human dorsolateral prefrontal cortex. Nature neuroscience, 24(3):425–436, 2021.

[4] Peng Xia, Juanhong Zhou, Rong Shen, and Degui Wang. Deciphering the cellular and molecular landscape of cervical cancer progression through single-cell and spatial transcriptomics. NPJ Precision Oncology, 9(1):158, 2025.

[5] Samuel G Rodriques, Robert R Stickels, Aleksandrina Goeva, Carly A Martin, Evan Murray, Charles R Vanderburg, Joshua Welch, Linlin M Chen, Fei Chen, and Evan Z Macosko. Slide-seq: A scalable technology for measuring genome-wide expression at high spatial resolution. Science, (6434):1463–1467, 2019.

[6] Kok Hao Chen, Alistair N Boettiger, Jeffrey R Moffitt, Siyuan Wang, and Xiaowei Zhuang. Spatially resolved, highly multiplexed rna profiling in single cells. Science, 348(6233):aaa6090, 2015.

[7] Lambda Moses and Lior Pachter. Museum of spatial transcriptomics. Nature Methods, 19(5):534–546, 2022.

[8] Ting Cui, Yan-Yu Li, Bing-Long Li, Han Zhang, Ting-Ting Yu, Jia-Ning Zhang, Feng-Cui Qian, Ming-Xue Yin, Qiao-Li Fang, Zi-Hao Hu, et al. Spatialref: a reference of spatial omics with known spot annotation. Nucleic Acids Research, 53(D1):D1215–D1223, 2025.

[9] Zhiyuan Yuan, Wentao Pan, Xuan Zhao, Fangyuan Zhao, Zhimeng Xu, Xiu Li, Yi Zhao, Michael Q Zhang, and Jianhua Yao. Sodb facilitates comprehensive exploration of spatial omics data. Nature Methods, 20(3):387–399, 2023.

[10] Shuangsang Fang, Bichao Chen, Yong Zhang, Haixi Sun, Longqi Liu, Shiping Liu, Yuxiang Li, and Xun Xu. Computational approaches and challenges in spatial transcriptomics. Genomics, Proteomics & Bioinformatics, 21(1):24–47, 2023.

[11] Ruben Dries, Qian Zhu, Rui Dong, Chee-Huat Linus Eng, Huipeng Li, Kan Liu, Yuntian Fu, Tianxiao Zhao, Arpan Sarkar, Feng Bao, et al. Giotto: a toolbox for integrative analysis and visualization of spatial expression data. Genome Biology, 22(1):78, 2021.

[12] Giovanni Palla, Hannah Spitzer, Michal Klein, David Fischer, Anna Christina Schaar, Louis Benedikt Kuemmerle, Sergei Rybakov, Ignacio L Ibarra, Olle Holmberg, Isaac Virshup, et al. Squidpy: a scalable framework for spatial omics analysis. Nature Methods, 19(2):171–178, 2022.

[13] Yihang Xiao, Jinyi Liu, Yan Zheng, Xiaohan Xie, Jianye Hao, Mingzhi Li, Ruitao Wang, Fei Ni, Yuxiao Li, Jintian Luo, et al. Cellagent: An llm-driven multi-agent framework for automated single-cell data analysis. In International Conference on Learning Representation, 2026.

[14] Nikita Mehandru, Amanda K Hall, Olesya Melnichenko, Yulia Dubinina, Daniel Tsirulnikov, David Bamman, Ahmed Alaa, Scott Saponas, and Venkat S Malladi. Bioagents: Bridging the gap in bioinformatics analysis with multi-agent systems. Scientific Reports, 15(1):39036, 2025.

[15] Qi Xin, Quyu Kong, Hongyi Ji, Yue Shen, Yuqi Liu, Yan Sun, Zhilin Zhang, Zhaorong Li, Xunlong Xia, Bing Deng, et al. Bioinformatics agent (bia): Unleashing the power of large language models to reshape bioinformatics workflow. BioRxiv, pages 2024–05, 2024.

[16] Yang Liu, Lu Zhou, Ruikun He, Rongbo Shen, and Yixue Li. Benchmarking llm-based agents for single-cell omics analysis. Genome Biology, 2026.

[17] Zhenyu Wang, Zikang Wang, Jiyue Jiang, Pengan Chen, Xiangyu Shi, and Yu Li. Large language models in bioinformatics: A survey. In Findings of the Association for Computational Linguistics, pages 3602–3615, 2025.

[18] Fnu Neha, Deepshikha Bhati, and Deepak Kumar Shukla. Retrieval-augmented generation (rag) in healthcare: A comprehensive review. AI, 6(9):226, 2025.

[19] Patrick Lewis, Ethan Perez, Aleksandra Piktus, Fabio Petroni, Vladimir Karpukhin, Naman Goyal, Heinrich Küttler, Mike Lewis, Wen-tau Yih, Tim Rocktäschel, et al. Retrieval-augmented generation for knowledge-intensive nlp tasks. Advances in neural information processing systems, 33:9459–9474, 2020.

[20] Florian Duclot and Mohamed Kabbaj. The role of early growth response 1 (egr1) in brain plasticity and neuropsychiatric disorders. Frontiers in Behavioral Neuroscience, 11:35, 2017.

[21] Tewarit Sarachana and Valerie W Hu. Genome-wide identification of transcriptional targets of rora reveals direct regulation of multiple genes associated with autism spectrum disorder. Molecular Autism, 4(1):14, 2013.

[22] Pierre J Magistretti and Igor Allaman. A cellular perspective on brain energy metabolism and functional imaging. Neuron, 86(4):883–901, 2015.

[23] Roshan Cools and Mark D’Esposito. Inverted-u–shaped dopamine actions on human working memory and cognitive control. Biological Psychiatry, 69(12):e113–e125, 2011.

[24] Christine Stadelmann, Sebastian Timmler, Alonso Barrantes-Freer, and Mikael Simons. Myelin in the central nervous system: structure, function, and pathology. Physiological Reviews, 2019.

[25] Jeremy K Seamans and Charles R Yang. The principal features and mechanisms of dopamine modulation in the prefrontal cortex. Progress in Neurobiology, 74(1):1–58, 2004.

[26] Hongkui Zeng, Elaine H Shen, John G Hohmann, Seung Wook Oh, Amy Bernard, Joshua J Royall, Katie J Glattfelder, Susan M Sunkin, John A Morris, Angela L Guillozet-Bongaarts, et al. Large-scale cellular-resolution gene profiling in human neocortex reveals species-specific molecular signatures. Cell, 149(2):483–496, 2012.

[27] Costantino Iadecola. The neurovascular unit coming of age: a journey through neurovascular coupling in health and disease. Neuron, 96(1):17–42, 2017.

[28] Amanda Janesick, Robert Shelansky, Andrew D Gottscho, Florian Wagner, Stephen R Williams, Morgane Rouault, Ghezal Beliakoff, Carolyn A Morrison, Michelli F Oliveira, Jordan T Sicherman, et al. High resolution mapping of the tumor microenvironment using integrated single-cell, spatial and in situ analysis. Nature communications, 14(1):8353, 2023.

[29] Quentin Blampey, Hakim Benkirane, Nadège Bercovici, Kevin Mulder, Grégoire Gessain, Florent Ginhoux, Fabrice André, and Paul-Henry Cournède. Novae: a graph-based foundation model for spatial transcriptomics data. Nature Methods, 22(12):2539–2550, 2025.

[30] Johanna A Joyce and Douglas T Fearon. T cell exclusion, immune privilege, and the tumor microenvironment. Science, 348(6230):74–80, 2015.

[31] Christian M Schürch, Salil S Bhate, Graham L Barlow, Darci J Phillips, Luca Noti, Inti Zlobec, Pauline Chu, Sarah Black, Janos Demeter, David R McIlwain, et al. Coordinated cellular neighborhoods orchestrate antitumoral immunity at the colorectal cancer invasive front. Cell, 182(5):1341–1359, 2020.

[32] Catherine Sautès-Fridman, Florent Petitprez, Julien Calderaro, and Wolf Herman Fridman. Tertiary lymphoid structures in the era of cancer immunotherapy. Nature Reviews Cancer, 19(6):307–325, 2019.

[33] Shunyu Yao, Jeffrey Zhao, Dian Yu, Nan Du, Izhak Shafran, Karthik R Narasimhan, and Yuan Cao. React: Synergizing reasoning and acting in language models. In The eleventh international conference on learning representations, 2022.

[34] Jeremy G Baldwin, Christoph Heuser-Loy, Tanmoy Saha, Roland C Schelker, Dragana Slavkovic-Lukic, Nicholas Strieder, Inmaculada Hernandez-Lopez, Nisha Rana, Markus Barden, Fabio Mastrogiovanni, et al. Intercellular nanotube-mediated mitochondrial transfer enhances t cell metabolic fitness and antitumor efficacy. Cell, 187(23):6614–6630, 2024.

[35] John Kwon and Samuel F Bakhoum. The cytosolic dna-sensing cgas–sting pathway in cancer. Cancer discovery, 10(1):26–39, 2020.

[36] Cameron G Williams, Hyun Jae Lee, Takahiro Asatsuma, Roser Vento-Tormo, and Ashraful Haque. An introduction to spatial transcriptomics for biomedical research. Genome Medicine, 14(1):68, 2022.

